# Functional regulatory evolution outside of the minimal *even-skipped* stripe 2 enhancer

**DOI:** 10.1101/101311

**Authors:** Justin Crocker, David L. Stern

**Affiliations:** Janelia Research Campus, Howard Hughes Medical Institute, 19700 Helix Drive, Ashburn, VA 20147 USA

**Keywords:** Evolution, Transcription, Enhancer, *Drosophila*, *Even-skipped*

## Abstract

Transcriptional enhancers are regions of DNA that drive gene expression at precise times, levels, and locations. While many studies have elucidated how individual enhancers can evolve, most of this work has focused on what are called “minimal” enhancers, the smallest DNA regions that drive expression that approximates an aspect of native gene expression. Here we explore how the *Drosophila erecta even-skipped* (eve) locus has evolved by testing its activity in the divergent *D. melanogaster* genome. We found, as has been reported previously, that the minimal *D. erecta eve* stripe 2 enhancer (*eveS2*) fails to drive appreciable expression in *D. melanogaster* [1]. However, we found that a large transgene carrying the entire *D. erecta eve* locus drives normal *eve* expression, including in stripe 2. We performed a functional dissection of the region upstream of the *D. erecta eveS2* region and found that regulatory information outside of the minimal *D. erecta eveS2* contains multiple Zelda motifs that are required for normal expression. Our results illustrate how sequences outside of minimal enhancer regions can evolve functionally through mechanisms other than changes in transcription factor binding sites that drive patterning.

## Introduction

Developmental enhancers contain multiple binding sites for transcription factors, together specifying the precise time, level, and location of gene expression. Classically, minimal enhancers have been identified as the smallest DNA fragments that are sufficient to direct reporter-gene expression in a particular tissue or domain of normal gene expression [2, 3]. These studies have provided mechanistic insight into how transcriptional logic is encoded in individual enhancers [4, 5]. However, in even the earliest studies, it was clear that minimal enhancers are insufficient to define the normal gene expression pattern with complete fidelity [2]. More recently, genomic studies have provided evidence that minimal enhancers are embedded within larger regions containing additional transcription factor binding sites that may be required for normal enhancer function [6, 7, 8].

Phenotypic evolution results largely from sequence changes in enhancers [9, 10, 11, 12, 13], even between closely related species [14, 15, 16, 17, 18, 19]. It is not clear, however, how often functional evolution includes changes within minimal enhancers versus outside of these regions. We have explored this problem through studies of the *even-skipped* (*eve*) gene.

The *Drosophila melanogaster eve* gene is expressed in seven transverse stripes along the anterior-posterior axis in the blastoderm embryo [20, 21]. Minimal enhancers have been identified that each direct expression in either one or two stripes that together drive expression in all seven stripes [22, 23]. Of all these enhancers, the minimal element for stripe 2 has been studied in the greatest detail [2, 24, 25]. This enhancer contains multiple binding sites for transcriptional activators (Bicoid and Hunchback) and repressors (Giant, Krüppel, and Sloppy-paired). The collective activity of transcription factors binding to these sites drives *eve* expression specifically in stripe 2 [2, 21, 24, 25, 26].

Previously, reporter gene assays were used to investigate the functional evolution of *eve S2* from three divergent *Drosophila* species with transgenic assays in *D. melanogaster* [1, 27, 15, 28, 6, 29]. These studies revealed that the *eveS2* enhancers from *D. yakuba* and *D. pseudoobscura*, which diverged ∼10 and ∼40 million years ago, respectively, from *D. melanogaster*, drove apparently normal expression in stripe 2. However, *eveS2* from *D. erecta,* which is closely related to *D. yakuba*, failed to drive appreciable levels of expression.

There are several possible reasons for why the *D. erecta eveS2* element does not drive appreciable expression in *D. melanogaster* [1]. First, the “minimal” functional *D. erecta eveS2* element may have been replaced elsewhere in the *D. erecta eve* locus with a functionally equivalent enhancer. Second, the *D. erecta* enhancer may contain all the required information to drive native expression, but the enhancer may have evolved to accommodate differences in the *D. erecta* embryonic environment [30, 31]—for example, differences in transcription factor concentrations [17, 32]. Third, the true functional *eveS2* enhancer may be larger than defined by the *D. melanogaster* minimal element and shifts in the locations of key transcription factor binding sites may have rendered the *D. erecta* region corresponding to the *D. melanogaster* “minimal” element unable to drive appropriate expression.

Here we tested these hypotheses with a functional dissection of the *D. erecta eve* locus in transgenic *D. melanogaster.* We found that a large transgene carrying the entire *D. erecta eve* locus drove normal expression in all seven stripes. This *D. erecta* transgene rescued down-stream *eve* targets and larval segmentation defects caused by an *eve* null mutation. We found that regulatory information required for *D. erecta* stripe 2 expression is located outside of the minimal stripe 2 enhancer region. Finally, we found that the *D. erecta* minimal *eveS2* region lacks multiple Zelda transcription factor binding sites that are found in *D. melanogaster* and we demonstrate that normal function of the *D. erecta eveS2* enhancer requires Zelda transcription factor binding motifs located outside of the minimal enhancer element. Zelda is apparently not required for patterning, but instead for making enhancers accessible for regulation [33, 34, 35, 36, 37, 38, 39, 40]. Many studies have suggested that the transcription factor binding sites required for normal enhancer function are often distributed over regions larger than experimentally-determined minimal enhancer elements and our results demonstrate that critical transcription factor binding sites can shift during evolution between locations within and outside of “minimal” regions.

## Results

*D. melanogaster* and *D. erecta* both express *eve* in seven embryonic stripes at similar levels and locations (Figure 1 a, b). A transgene of the *D. melanogaster eveS2* minimal enhancer, when re-introduced into *D. melanogaster,* drives robust expression in approximately the same region as the native expression of *eve* stripe 2 (Figure 1c). However, the orthologous *eve S2* fragment from *D. erecta* does not drive expression in transgenic *D. melanogaster* (Figure 1d) (see also Ludwig et al., [1]). A 20kb region of the *D. melanogaster* genome surrounding the *eve* transcription unit is sufficient to drive apparently normal *eve* expression and rescues transcription of genes that are normally regulated by Eve [23]. To maximize the likelihood that we would capture the entire *eve* locus from *D. erecta,* we tested the ability of a ~47 kb *D. erecta* fosmid to drive expression in *D. melanogaster* embryos deficient for native *eve* function. This fosmid contains *D. erecta* sequences orthologous to all of the *D. melanogaster* stripe enhancers (Fig. 1e).

**Figure 1.**
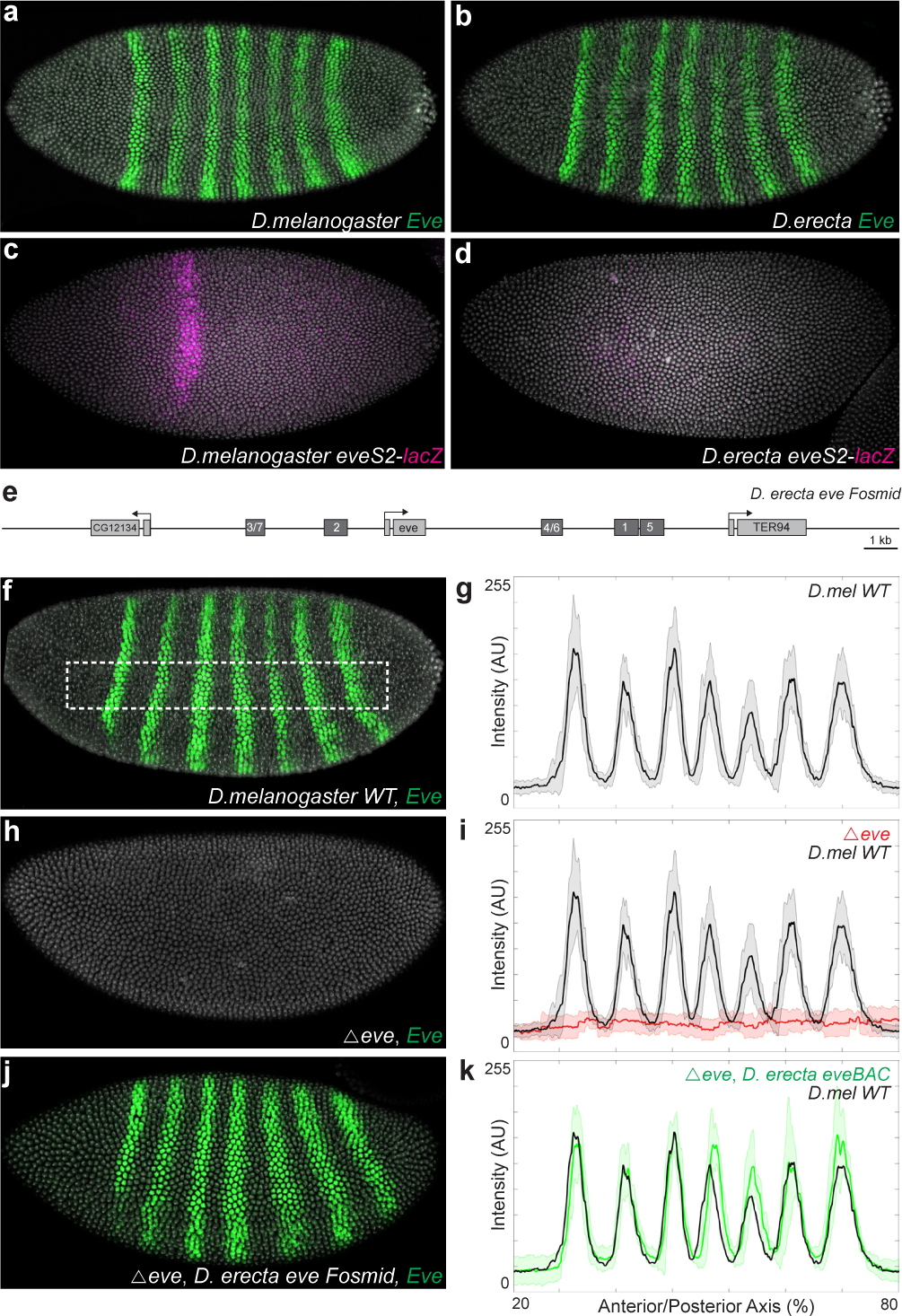
The *D. erecta eve* locus drives normal *eve* expression in transgenic *D. melanogaster.* (**a, b**) Stage 5 embryos stained for Eve protein in either *D. melanogaster* (**a**) or *D. erecta* (**b**). (**c**, **d**) Stage 5 *D. melanogaster* embryos stained for *ß-Gal* RNA carrying the D. *melanogaster eveS2* (**c**) or *D. erecta eveS2.* **e**) Schematic representation of the *eve* locus with enhancers indicated in dark boxes. (**f**, **h**, **j**) Stage 5 embryos stained for Eve protein in either a wild-type background (**f**), *eve* null background (h), or an *eve* null background carrying the *D. erecta eve* fosmid (**j**). (**g**, **i**, **k**) Profiles of average expression levels across the region indicated in panel (**f**) for the indicated genotype (n=10 for each genotype). In all plots, the solid black line denotes wild-type, red denotes eve null (**i**), and green denotes an *eve* null background carrying the *D. erecta eve* fosmid (**k**). Bounding areas around experimental data indicate one standard deviation. AU indicates Arbitrary Units of fluorescence intensity.

We found that this *D. erecta* fosmid drove apparently normal expression of all seven *eve* stripes (Fig. 1f-k). To determine whether the timing, levels, and spatial distribution of Eve expression driven by the *D. erecta* fosmid were correct in *D. melanogaster,* we tested the ability of the *D. erecta* locus to functionally rescue gene regulatory networks that are normally regulated by Eve. We found that the *D. erecta eve* fosmid rescued the expression of the segmental polarity gene *engrailed* (Fig. 2a-c). Additionally, we found that the fosmid rescued all segmentation defects in the cuticles of larvae deficient for *eve* (Fig. 2d-f). Together, these results suggest that the 47kb *D. erecta* fosmid contains all of the regulatory information required for *eve* expression in stripe 2 and all other *eve* stripes.

**Figure 2.**
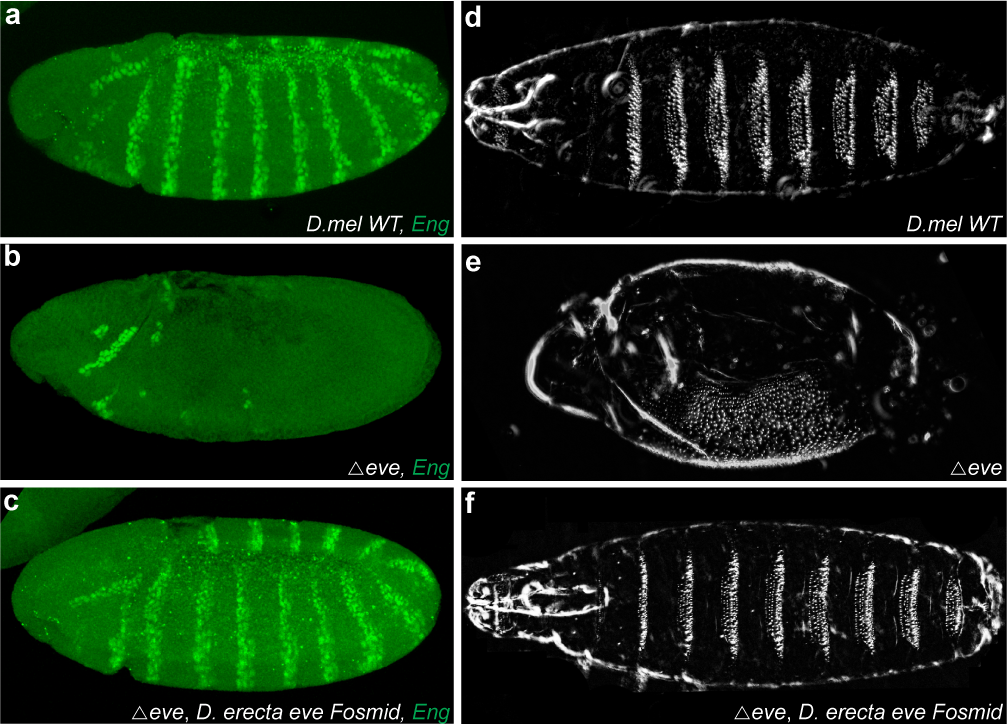
Functional rescue of eve null flies by the D. erecta fosmid. (**a, b, c**) Stage 9 embryos stained for Engrailed (Eng) protein in either a wild-type background (**a**), eve null background (**b**), or an eve null background carrying the D. erecta eve fosmid (**c**). (**d, e, f**) First instar larval cuticle preps of either a wild-type (**d**), eve null background (**e**), or an eve null background carrying the D. erecta eve fosmid (**f**).

The regulatory information for *D. melanogaster eve* stripe 2 and stripes 3/7 is located upstream of the *eve* promoter [41, 2, 25, 42]. Increasing the size of the DNA regions tested for these enhancers increases levels of transcription [41, 2, 25, 42]. Additionally, while the minimal *D. melanogaster* enhancer is sufficient to drive approximately normal expression of *eve* stripe 2, sequences surrounding the minimal *eveS2* element contribute to the robustness of the minimal enhancer [6]. Therefore, we suspected that sequences near the minimal *D. erecta eveS2* may be required for normal expression.

To test if regulatory information 5’ of the minimal *D. erecta eveS2* region is required for normal stripe 2 expression, we tested a series of constructs that included the minimal stripe 2 enhancer plus progressively more 5’ DNA (Fig. 3a). While the 855 bp *D. erecta* minimal *eveS2* construct drove very little expression in *D. melanogaster* (Fig. 3b, c), a fragment containing an additional 832 bp upstream of the minimal enhancer drove weak expression in the stripe two domain (Fig. 3 d, e). Increasing the minimal construct size with an additional 1609 bp 5’ of the minimal enhancer, up to the boundary with the minimal stripe 3/7 enhancer, further increased levels of stripe 2 expression and drove weak expression in stripes 3 and 7 (Fig. 3 f, g). This indicates both that information critical for stripe 2 expression resides outside the minimal stripe 2 region in *D. erecta*, but also that patterning information for stripes 3 and 7 resides outside of the minimal stripe 3/7 enhancer. Increasing the size of the element further, so that it encompassed the minimal *eve3/7* enhancer, did not further increase expression levels in the domain of *eve* stripe two, but increased expression in stripes 3 and 7 (Fig. 3h, i). Together, these results demonstrate that up to 1609 bp 5’ of the minimal *D. erecta eve* stripe-two enhancer contains regulatory information required for *eve* expression in transgenic *D. melanogaster*.

**Figure 3.**
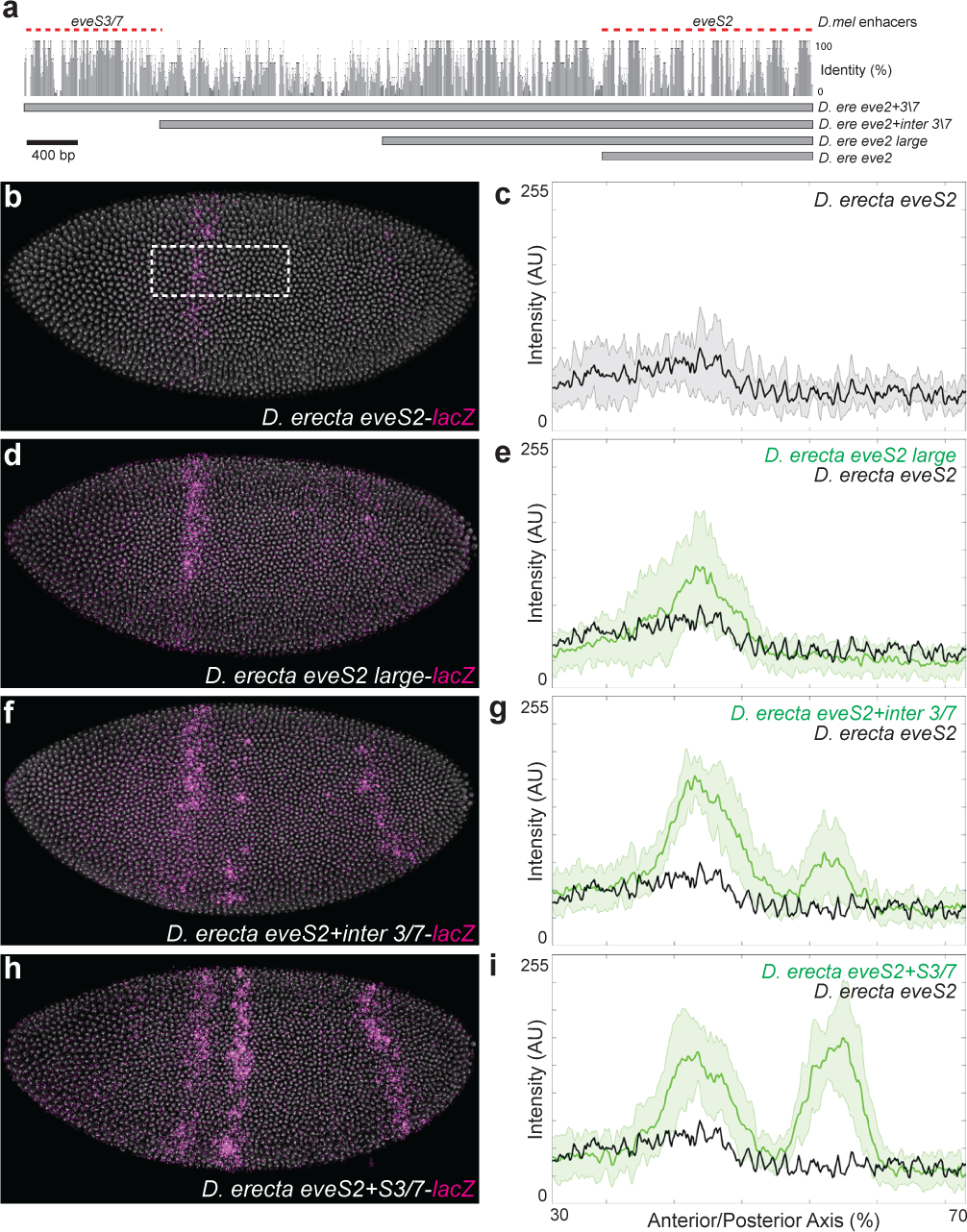
Sequences upstream of the minimal *D. erecta eveS2* enhancer drive expression in *D. melanogaster*. (**a**) Schematic representation of the *eve* locus with enhancers tested indicated. The sequence conservation plot is a 5-way sequence alignment between *D. melanogaster, sechellia, yakuba, erecta*, and *ananassae*. (**b**, **d**, **f**, **h**) Stage 5 *D. melanogaster* embryos stained for *ß-Gal* RNA carrying the indicated transgene. (**c**, **e**, **g**, **i**) Profiles of average expression levels across the region indicated in panel (**b**) for the indicated enhancer (n=10 for each genotype). In all plots, the solid black line denotes wild-type, and green denotes the *D. erecta eve* enhancer (**k**). Bounding areas around experimental data indicate one standard deviation. AU indicates Arbitrary Units of fluorescence intensity.

We next performed a computational search for binding sites of three transcription factors that regulate the spatial pattern of the *D. melanogaster* eveS2 enhancer—Bicoid, Hunch-back and Krüppel—across the regulatory region upstream of the minimal enhancer (Fig. S1). We observed no obvious turnover of these binding sites in *D. erecta* that would explain the loss of expression from the minimal *eveS2* enhancer, consistent with previous studies [1].

We next examined the spatial distribution of putative binding sites for the Zelda protein. *Zelda* is expressed ubiquitously in the blastoderm embryo and Zelda protein binds to many enhancers that drive transcription in the early blastoderm embryo [33, 34, 35, 36, 37, 38, 39, 40]. Zelda activity is correlated with chromatin accessibility [35, 39, 40] and Zelda appears to make enhancers accessible to transcription factors that drive specific patterns of gene expression [34, 35, 38, 39, 40, 43].

We searched for putative Zelda motifs in the eveS2 minimal element in *D. melanogaster, D. sechellia, D. yakuba*, and *D. erecta*. We found four sites that were perfectly conserved between these species, and six sites that were present in *D. melanogaster* but absent in *D. erecta.* It is possible that the loss of these motifs has led to the loss of activity of the *D. erecta eveS2* minimal element. Because the larger *D. erecta* reporter constructs we tested drove stripe 2 expression, we searched for additional Zelda motifs 5’ of the minimal element. We identified three Zelda motifs in this region in *D. erecta,* two of which are conserved in *D. melanogaster* and one of which is absent in *D. melanogaster* (Fig. 4a, b). We tested the activity of these motifs *in vivo* by deleting all three motifs from the large *D. erecta eveS2* construct (Fig. 4c, d). Removal of these upstream Zelda motifs abrogated reporter gene expression (Fig. 4d). These Zelda motifs are therefore critical for stripe 2 expression in *D. erecta*.

**Figure 4.**
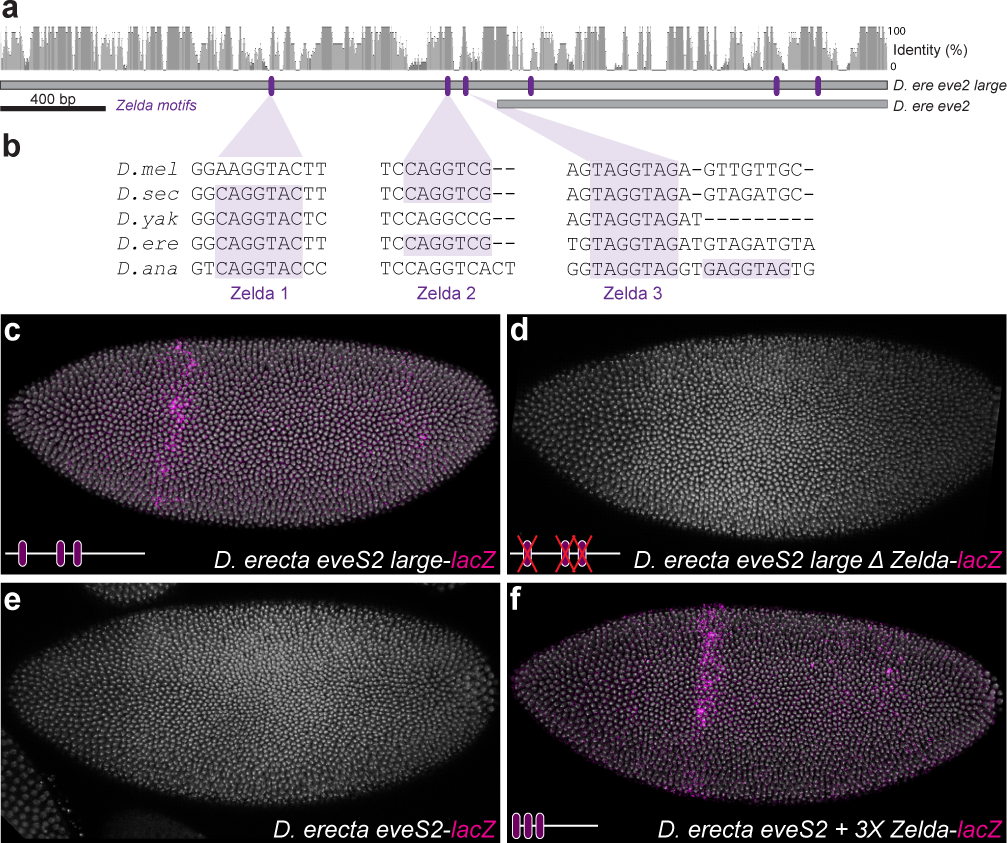
Zelda motifs are necessary and sufficient for the expression of the minimal *D. erecta eveS2*. (**a**, **b**) Schematic representation of the *eve* locus, with tested enhancers indicated, highlighting the upstream Zelda motifs in *D. erecta* (**b**). The sequence conservation plot is a 5-way sequence alignment between *D. melanogaster, sechellia, yakuba, erecta,* and *ananassae.* (**c**, **d**, **e**, **f**) Stage 5 *D. melanogaster* embryos stained for *ß-Gal* RNA carrying the indicated *D. erecta* transgene

To test if these Zelda motifs were sufficient to drive expression when appended to the minimal *D. erecta eveS2*, we placed the three Zelda motifs upstream of the minimal element. Strikingly, we found that this construct drove expression in stripe 2 (Fig. 4e, f). Addition of these Zelda sites is therefore sufficient to allow properly patterned expression of the minimal *D. erecta* stripe 2 element. Furthermore, the patterns of Zelda motif gain and loss suggest that loss of Zelda motifs within the minimal elements prevents the *D. erecta* minimal enhancer element from driving expression and that gain of a new Zelda motif upstream of the minimal element may be required for normal *D. erecta* stripe 2 expression.

## Discussion

Our results suggest that functional regulatory evolution has occurred outside the minimal *eveS2* enhancer between *D. melanogaster* and *D. erecta.* The minimal *D. erecta eveS2* region contains binding sites necessary for patterning of expression in stripe 2, but lacks sufficient Zelda binding sites to drive expression. These results support the hypothesis that Zelda contributes to determining the regulatory state of the *eveS2* enhancer in the blastoderm embryo, ON versus OFF, and can be decoupled, at least in part, from the patterns and levels of expression driven by the enhancer [34, 35, 38, 39, 40, 43]. We have not explored the 777 bp upstream of these Zelda sites (the extra DNA in construct *D. ere eve 2+inter 3\7*) for the sequences that drive stronger expression in stripe 2. This is a poorly conserved genomic region and it is curious that it contains information both for bolstering stripe 2 expression and for patterning stripes 3 and 7. The function of this DNA region would repay further exploration.

This study highlights the need to understand gene regulatory evolution at the level of the entire locus. While studies of minimal enhancers elements have provided enormous insight into the mechanisms of gene regulation, the broader DNA regions around minimal enhancers also contribute to enhancer function and evolution [44, 45, 46, 47, 48, 49, 50]. Our results encourage great caution in comparative studies of enhancer evolution when functional studies have been performed on enhancers from only one species. One goal for future research is to understand how gene regulation evolves at the level of the entire locus, including information such as binding site composition, the arrangement of enhancers, the activities of intervening sequences, and the distance between functional sequences.

## Methods and Materials

### 0.1 Construction of enhancer constructs

The *D. erecta* locus was tested by cloning the BDERF01-4213 fosmid, obtained from BACPAC resources (https://bacpacresources.org/), corresponding to *D. melanogaster* chr2R:5,840,325-5,887,364, into vector TKBL-w+ [51]. All enhancer constructs were cloned into the placZattB expression construct with a hsp70 promoter [18]. See supplemental materials for complete construct sequences.

### 0.2 Fly strains and crosses

*D. melanogaster* strains were maintained under standard laboratory conditions. Transgenic enhancer constructs were created by Rainbow Transgenic Flies Inc. and were integrated at the attP2 landing site. The *eveR13* mutation was used to test for complementation using the *D. erecta* fosmid. The lethal mutations were balanced over marked balancer chromosome *CyO P(hb-lacZ)* to allow identification of mutant embryos by immunostaining for Beta-galactosidase.

### 0.3 Embryo manipulations

Embryos were raised at 25° C and fixed and stained according to standard protocols [52]. Briefly, primary antibodies obtained from the Developmental Studies Hybridoma Bank were used to detect Eve (3C10, used 1:20) and En (4D9, used 1:20) proteins, which was followed by detection of primary antibodies using secondary antibodies labeled with Alexa Fluor dyes (1:500, Invitrogen). Cuticle preps were performed using standard protocols [53].

### 0.4 Microscopy

Each series of experiments to measure transcript levels was performed entirely in parallel. Embryo collections, fixations, and hybridizations, image acquisitions, and processing were performed side-by-side in identical conditions. Confocal exposures were identical for each series and were set to not exceed the 255 maximum level. Confocal images were obtained on a Leica DM5500 Q Microscope with an ACS APO 20×/0.60 IMM CORR lens and Leica Microsystems LAS AP software. Sum projections of confocal stacks were assembled, embryos were scaled to match sizes, background was subtracted using a 50-pixel rolling-ball radius and plot profiles of fluorescence intensity were analyzed using ImageJ software (http://rsb.info.nih.gov/ij/). Data from the plot profiles were further analyzed in Matlab.

## Acknowledgments

We thank the entire Stern lab for discussion and comments, Claire Standley for comments on the manuscript, and Jessica Cande for the Bourbon fruit cake that proved important for manuscript preparation.

## Appendices

### Supplemental Figures

**Figure S1.**
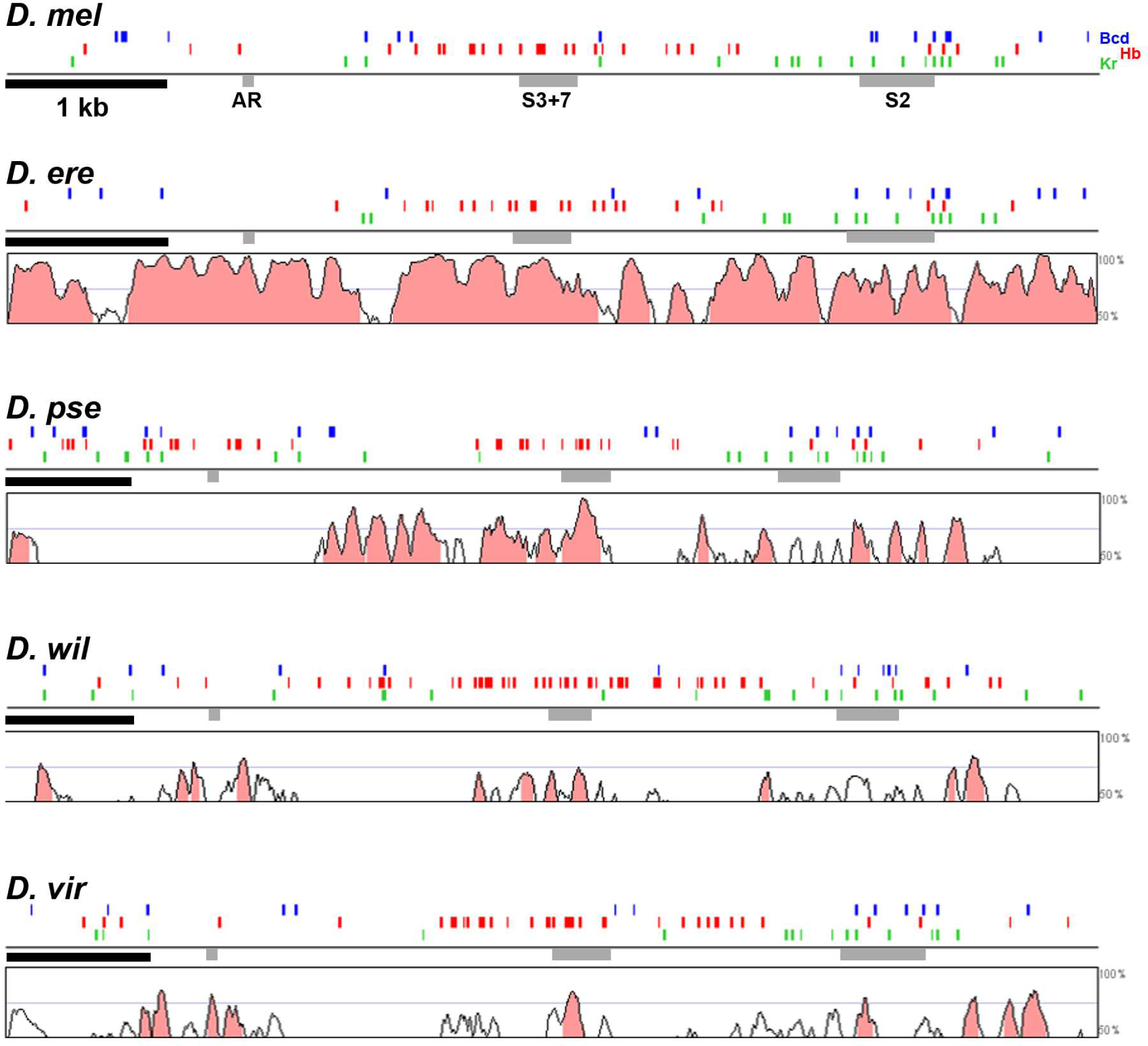
VISTA plot demonstrating extensive sequence conservation upstream of the eve start site [54]. Gray boxes denote regions homologous to the *D. melanogaster* minimal regulatory elements. Computationally predicted binding sites based on reported position-weighted matrices [55], demonstrate an extensive number of conserved binding sites for Bcd (Blue), Hb (Red), and Kr (Green)

**Figure S2.**
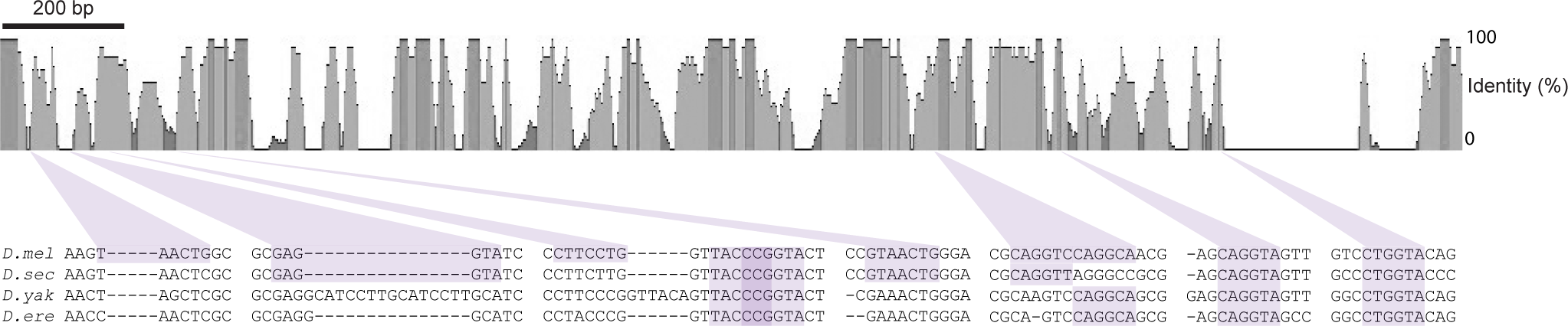
Multiple species alignment of the *eveS2* minimal enhancer sequence with Zelda motifs and conservation noted. Highlighted Zelda motifs denote the minimal binding site sequence 5’-CAGGTA [40], allowing for one mismatch to identify possible low affinity Zelda binding sites [56, 18].

## Supplemental Construct Sequences

>D. erecta eveS2

CGGCGCAATATAACCCAATAATTTGAACCAACTCGCGGAGCAGCGAGGGCATCCTACCCGGTTACCCGGTACTGCATAACAATGAAACGAAACTGGGACAGATCGGTGATGGTTTCTCGCTGTGTGTGCCGTGTTAATCCGTTTGCCATCAGCCAGATTATTAGTCAATTGCAGTCGCAGTCGCAGTTGCAGTTGCAGGGTTTCGCTTTCCTCGTCCTCGTTTCACTTTCGAGTTAGACTTTATTGCAGCATCTTGCAGCAACAATCGGCGCAGTTTGGTAACACGCTGTGCCCTTCCACTTTCCACATTCCACGGCCCAATTCGGCGGATTTAGACGGAATCGAGGGACCCTGACTATGTTCGCATAATGAAAGCGAAACCAAACCGGGTTGCAAAGTCAGGGCATTCCATCGCCGTCGCCATCGCCACCGCCATCTTCTGCGGGCGTTTGTTTGTTTGTTTGCTGGGATTAGCCAAGGGCTTGACTTGGAACCCAATCCCAATCCCAATCCCAATCCCAAAGCCAATCCCAGTGCCCATCCCGATCGCAGTCCCATGCCCTTTTCATTAGAAAGTCATAAAAACACATAATAATGATGTCGAACGGATTAGCTGCGCGCAGTCCAGGCAGCGCAATTAACGGACTAGCAAACTGGGTTATTTCTATTTTTTTATTTTCGCCGACTTAGCCCTGATCCGCGAGCTTAACCCGTTTGAGCCGCAGCAGGTAGCCATCCCCATCCTGGCTTGCGCAAGTGGCAGGTTCGCTGCCCCAGGAGAGCCGTGGAGGCACTCTGGCCACTGGCCTGGTACAGTTGCCGCTGGGCATGATTATATCATCATAATAAATGTTT

>D. erecta eveS2 large

TCCTCGAGGATTTCGGGCTGAATCACTTACTCAACCCGTCGATTCCGTCCGCGCAATCATCATAAATTCTCGGTCTTTTTGCTGTAATTGTTTTATGGCAGAAATTACTCAATCATCAAGCATAATTCCCTCGTTTTCGCCGTTTTATTGCCAATTTTTGCACTGCCTCCCGCCTTTGCCACTCCCGGCCCTTCCCTCATCGTTTTGCGAATCTCCGACGGATTCGCATTTCTATTGCGCGGACAATCCGGCCAGTGTGTTTGCCATTTACTTGCCATGATGACGGGCATAATCAGCGAGATCGGCGCTTTGTGAGTGCAGAATGTGCAATAAAGCGGCAACAATCGGCTTGGGATTCGCCTTCCCCTATTCCCAGTATTGCCCGAGTGCCCGGACGACTGCGAAAGTGTTTGCGGATCGGGATCGGAATCGGAATAGGAATAGGAATACGAGACTGAGCAGAGGCAGGTACTTCCCGCCGGCCGGACACTTTCGCCTAACCAAGCGGACTCAACCCAACCCAACCCAACACAACCCAATCCAACCCACCCGATCGCCATAAAGGGTATTTACTGTCGCTGCCGCAGAGCCTCGCTTGACGACTTAACCCAAGCGGTCGTTCCACGTCCATTCTCCGGACGGAGTCAAAGACAAAGGCCGGCGGAGCTGGACAATAGGCAAGGTTGTTGCTTGTGGGTAGAGGGTTTCAATCCCGAAATTGTACCTTTCCCGGGCGAGAAGGGCTTGCATGTGGGCCTTTTCCAGGTCGCAGATGTAGGTAGATGTAGATGTAGATGCGGATGGGCCGGTCGAGTTAATGCCAATGCAAATTGCGGCGCAATATAACCCAATAATTTGAACCAACTCGCGGAGCAGCGAGGGCATCCTACCCGGTTACCCGGTACTGCATAACAATGAAACGAAACTGGGACAGATCGGTGATGGTTTCTCGCTGTGTGTGCCGTGTTAATCCGTTTGCCATCAGCCAGATTATTAGTCAATTGCAGTCGCAGTCGCAGTTGCAGTTGCAGGGTTTCGCTTTCCTCGTCCTCGTTTCACTTTCGAGTTAGACTTTATTGCAGCATCTTGCAGCAACAATCGGCGCAGTTTGGTAACACGCTGTGCCCTTCCACTTTCCACATTCCACGGCCCAATTCGGCGGATTTAGACGGAATCGAGGGACCCTGACTATGTTCGCATAATGAAAGCGAAACCAAACCGGGTTGCAAAGTCAGGGCATTCCATCGCCGTCGCCATCGCCACCGCCATCTTCTGCGGGCGTTTGTTTGTTTGTTTGCTGGGATTAGCCAAGGGCTTGACTTGGAACCCAATCCCAATCCCAATCCCAATCCCAAAGCCAATCCCAGTGCCCATCCCGATCGCAGTCCCATGCCCTTTTCATTAGAAAGTCATAAAAACACATAATAATGATGTCGAACGGATTAGCTGCGCGCAGTCCAGGCAGCGCAATTAACGGACTAGCAAACTGGGTTATTTCTATTTTTTTATTTTCGCCGACTTAGCCCTGATCCGCGAGCTTAACCCGTTTGAGCCGCAGCAGGTAGCCATCCCCATCCTGGCTTGCGCAAGTGGCAGGTTCGCTGCCCCAGGAGAGCCGTGGAGGCACTCTGGCCACTGGCCTGGTACAGTTGCCGCTGGGCATGATTATATCATCATAATAAATGTTT

>D. erecta eveS2+inter 3/7

TTCGTTCCCAACTACGGCTAAGATATGCCAGTTTGTTTTGTCTCCGGCAATTATTGGAAATTTCATTGGGTCGATTGGGTAGATGTCGATTGGGTCGATGTCGATGCCTTCCCTCGGGAAAAGTGAATAGGTTGTGCCATAAAAATCGCTGCTCTTGGAGATGAAATGCTGTAGTAGTATGCCAAAGCATCATTCTGCTTTTTTATTTTCTCACTGCTAAATGCAGCTAATTTGTCGATTGTCTGAAAAGTGTTCTCTAAGCCGAAGCACTTTTTTATGATGTTGCTAGAAAATGAAATCACTTATGTTGCCATACATCCCCAGGCATTTTATGGCCATTTGAGTGCGGGGTGCGCAGTTCTGCTTAAGTGGCGGATGGAAACCACCACATTTATTCGAGGGATGATGTGCTCTAATACCTCCTCATCAAATGGGATGGTCTCTTCGCATGGAGAGTGGCAAACTCTTGGAAAAGTGAGGCGGAGTTAAAAAGCCGTGCTATATAACAAGATTTTTGATATTCAGTTATGTATATATGGACAAGAAATATCAAAGACCTTATCAAATATGTTGCCTTTTATCCTCGAAATGAAACAAATGCTCTACTAATTTGGCAAGTCAAAAGATCAGTTCAGCAATTATTCAAAAGAACATAAAATATGCGTATATTTTTGGGAATGTACCAGTGCTTTCCAAAATAGATTGCCAAACAAATCAACTAATAACTTTAATTTAAAAAACTGGGCAATCCTGAGTTGGCAGTCTTCCCAAGAATGGCTCCTCGAGGATTTCGGGCTGAATCACTTACTCAACCCGTCGATTCCGTCCGCGCAATCATCATAAATTCTCGGTCTTTTTGCTGTAATTGTTTTATGGCAGAAATTACTCAATCATCAAGCATAATTCCCTCGTTTTCGCCGTTTTATTGCCAATTTTTGCACTGCCTCCCGCCTTTGCCACTCCCGGCCCTTCCCTCATCGTTTTGCGAATCTCCGACGGATTCGCATTTCTATTGCGCGGACAATCCGGCCAGTGTGTTTGCCATTTACTTGCCATGATGACGGGCATAATCAGCGAGATCGGCGCTTTGTGAGTGCAGAATGTGCAATAAAGCGGCAACAATCGGCTTGGGATTCGCCTTCCCCTATTCCCAGTATTGCCCGAGTGCCCGGACGACTGCGAAAGTGTTTGCGGATCGGGATCGGAATCGGAATAGGAATAGGAATACGAGACTGAGCAGAGGCAGGTACTTCCCGCCGGCCGGACACTTTCGCCTAACCAAGCGGACTCAACCCAACCCAACCCAACACAACCCAATCCAACCCACCCGATCGCCATAAAGGGTATTTACTGTCGCTGCCGCAGAGCCTCGCTTGACGACTTAACCCAAGCGGTCGTTCCACGTCCATTCTCCGGACGGAGTCAAAGACAAAGGCCGGCGGAGCTGGACAATAGGCAAGGTTGTTGCTTGTGGGTAGAGGGTTTCAATCCCGAAATTGTACCTTTCCCGGGCGAGAAGGGCTTGCATGTGGGCCTTTTCCAGGTCGCAGATGTAGGTAGATGTAGATGTAGATGCGGATGGGCCGGTCGAGTTAATGCCAATGCAAATTGCGGCGCAATATAACCCAATAATTTGAACCAACTCGCGGAGCAGCGAGGGCATCCTACCCGGTTACCCGGTACTGCATAACAATGAAACGAAACTGGGACAGATCGGTGATGGTTTCTCGCTGTGTGTGCCGTGTTAATCCGTTTGCCATCAGCCAGATTATTAGTCAATTGCAGTCGCAGTCGCAGTTGCAGTTGCAGGGTTTCGCTTTCCTCGTCCTCGTTTCACTTTCGAGTTAGACTTTATTGCAGCATCTTGCAGCAACAATCGGCGCAGTTTGGTAACACGCTGTGCCCTTCCACTTTCCACATTCCACGGCCCAATTCGGCGGATTTAGACGGAATCGAGGGACCCTGACTATGTTCGCATAATGAAAGCGAAACCAAACCGGGTTGCAAAGTCAGGGCATTCCATCGCCGTCGCCATCGCCACCGCCATCTTCTGCGGGCGTTTGTTTGTTTGTTTGCTGGGATTAGCCAAGGGCTTGACTTGGAACCCAATCCCAATCCCAATCCCAATCCCAAAGCCAATCCCAGTGCCCATCCCGATCGCAGTCCCATGCCCTTTTCATTAGAAAGTCATAAAAACACATAATAATGATGTCGAACGGATTAGCTGCGCGCAGTCCAGGCAGCGCAATTAACGGACTAGCAAACTGGGTTATTTCTATTTTTTTATTTTCGCCGACTTAGCCCTGATCCGCGAGCTTAACCCGTTTGAGCCGCAGCAGGTAGCCATCCCCATCCTGGCTTGCGCAAGTGGCAGGTTCGCTGCCCCAGGAGAGCCGTGGAGGCACTCTGGCCACTGGCCTGGTACAGTTGCCGCTGGGCATGATTATATCATCATAATAAATG

>D. erecta eveS2+S3/7

GGACACAAGGATCCTCGAAATCGAGAGCGACCTCGCTGCATTAGAAAACTAGATCAGTTTTTTGTTTCGCGTCCGCTGATTTTTGTGCCCGTTTGCTCTCTTTACGGTTTATGGCCCCGTTTCCATTATTTCGTCATTTTTCCACATTTCCCAGCTCCTTTGTGCCGCTCAAAGAAATCTGTACGGAATTATGGTATATGCAGATTTTTATGGGTCGCCGATCCGGTTCGCGGAACGCGAGTGTCCTGCCGCGAGGACCTCAGCGGCGATCCTTGTCGGCCGTATTAAGAAAGTAGATCACGTTTTTTGTTCCCATTGTGCGCTTTTTTCGCTGCGCCAGTTCTGTTCCCCGAACCCAGCGAACTGCTCTAATTTTTTAATTCTTCACGGACTTTCATTGGGCTCCTGGAAAAACACGCACAAGGCTAGCTCTAGGACTCTACTGGCATACCTGTAATGGTGTCCATAACGCGCACTGCTCTCGTTTTTAAGACCCGTTTGTGTTCGTCTCTGGAAAGTAGATGGCATTCACATTTTTATGAGTTCGTTCCCAACTACGGCTAAGATATGCCAGTTTGTTTTGTCTCCGGCAATTATTGGAAATTTCATTGGGTCGATTGGGTAGATGTCGATTGGGTCGATGTCGATGCCTTCCCTCGGGAAAAGTGAATAGGTTGTGCCATAAAAATCGCTGCTCTTGGAGATGAAATGCTGTAGTAGTATGCCAAAGCATCATTCTGCTTTTTTATTTTCTCACTGCTAAATGCAGCTAATTTGTCGATTGTCTGAAAAGTGTTCTCTAAGCCGAAGCACTTTTTTATGATGTTGCTAGAAAATGAAATCACTTATGTTGCCATACATCCCCAGGCATTTTATGGCCATTTGAGTGCGGGGTGCGCAGTTCTGCTTAAGTGGCGGATGGAAACCACCACATTTATTCGAGGGATGATGTGCTCTAATACCTCCTCATCAAATGGGATGGTCTCTTCGCATGGAGAGTGGCAAACTCTTGGAAAAGTGAGGCGGAGTTAAACAAAAAGTCACTGGTTGTCGTCACTTATTTATAACTACACGTACGTTATATATTGGAATAAATCTACAAGATTTGACTTGCTTAATCATATAATTGTTGGAGCCAAAAGGATACAATAAATACGAAATACGAAGCTCTTCTAAGTCAATTATTCAAAAGAACATAAAATATGCGTATATTTTTGGGAATGTACCAGTGCTTTCCAAAATAGATTGCCAAACAAATCAACTAATAACTTTAATTTAAAAAACTGGGCAATCCTGAGTTGGCAGTCTTCCCAAGAATGGCTCCTCGAGGATTTCGGGCTGAATCACTTACTCAACCCGTCGATTCCGTCCGCGCAATCATCATAAATTCTCGGTCTTTTTGCTGTAATTGTTTTATGGCAGAAATTACTCAATCATCAAGCATAATTCCCTCGTTTTCGCCGTTTTATTGCCAATTTTTGCACTGCCTCCCGCCTTTGCCACTCCCGGCCCTTCCCTCATCGTTTTGCGAATCTCCGACGGATTCGCATTTCTATTGCGCGGACAATCCGGCCAGTGTGTTTGCCATTTACTTGCCATGATGACGGGCATAATCAGCGAGATCGGCGCTTTGTGAGTGCAGAATGTGCAATAAAGCGGCAACAATCGGCTTGGGATTCGCCTTCCCCTATTCCCAGTATTGCCCGAGTGCCCGGACGACTGCGAAAGTGTTTGCGGATCGGGATCGGAATCGGAATAGGAATAGGAATACGAGACTGAGCAGAGGCAGGTACTTCCCGCCGGCCGGACACTTTCGCCTAACCAAGCGGACTCAACCCAACCCAACCCAACACAACCCAATCCAACCCACCCGATCGCCATAAAGGGTATTTACTGTCGCTGCCGCAGAGCCTCGCTTGACGACTTAACCCAAGCGGTCGTTCCACGTCCATTCTCCGGACGGAGTCAAAGACAAAGGCCGGCGGAGCTGGACAATAGGCAAGGTTGTTGCTTGTGGGTAGAGGGTTTCAATCCCGAAATTGTACCTTTCCCGGGCGAGAAGGGCTTGCATGTGGGCCTTTTCCAGGTCGCAGATGTAGGTAGATGTAGATGTAGATGCGGATGGGCCGGTCGAGTTAATGCCAATGCAAATTGCGGCGCAATATAACCCAATAATTTGAACCAACTCGCGGAGCAGCGAGGGCATCCTACCCGGTTACCCGGTACTGCATAACAATGAAACGAAACTGGGACAGATCGGTGATGGTTTCTCGCTGTGTGTGCCGTGTTAATCCGTTTGCCATCAGCCAGATTATTAGTCAATTGCAGTCGCAGTCGCAGTTGCAGTTGCAGGGTTTCGCTTTCCTCGTCCTCGTTTCACTTTCGAGTTAGACTTTATTGCAGCATCTTGCAGCAACAATCGGCGCAGTTTGGTAACACGCTGTGCCCTTCCACTTTCCACATTCCACGGCCCAATTCGGCGGATTTAGACGGAATCGAGGGACCCTGACTATGTTCGCATAATGAAAGCGAAACCAAACCGGGTTGCAAAGTCAGGGCATTCCATCGCCGTCGCCATCGCCACCGCCATCTTCTGCGGGCGTTTGTTTGTTTGTTTGCTGGGATTAGCCAAGGGCTTGACTTGGAACCCAATCCCAATCCCAATCCCAATCCCAAAGCCAATCCCAGTGCCCATCCCGATCGCAGTCCCATGCCCTTTTCATTAGAAAGTCATAAAAACACATAATAATGATGTCGAACGGATTAGCTGCGCGCAGTCCAGGCAGCGCAATTAACGGACTAGCAAACTGGGTTATTTCTATTTTTTTATTTTCGCCGACTTAGCCCTGATCCGCGAGCTTAACCCGTTTGAGCCGCAGCAGGTAGCCATCCCCATCCTGGCTTGCGCAAGTGGCAGGTTCGCTGCCCCAGGAGAGCCGTGGAGGCACTCTGGCCACTGGCCTGGTACAGTTGCCGCTGGGCATGATTATATCATCATAATAAATGTTT

>D. erecta eveS2 large del. Zelda

TCCTCGAGGATTTCGGGCTGAATCACTTACTCAACCCGTCGATTCCGTCCGCGCAATCATCATAAATTCTCGGTCTTTTTGCTGTAATTGTTTTATGGCAGAAATTACTCAATCATCAAGCATAATTCCCTCGTTTTCGCCGTTTTATTGCCAATTTTTGCACTGCCTCCCGCCTTTGCCACTCCCGGCCCTTCCCTCATCGTTTTGCGAATCTCCGACGGATTCGCATTTCTATTGCGCGGACAATCCGGCCAGTGTGTTTGCCATTTACTTGCCATGATGACGGGCATAATCAGCGAGATCGGCGCTTTGTGAGTGCAGAATGTGCAATAAAGCGGCAACAATCGGCTTGGGATTCGCCTTCCCCTATTCCCAGTATTGCCCGAGTGCCCGGACGACTGCGAAAGTGTTTGCGGATCGGGATCGGAATCGGAATAGGAATAGGAATACGAGACTGAGCAGAGGAATTTACTTCCCGCCGGCCGGACACTTTCGCCTAACCAAGCGGACTCAACCCAACCCAACCCAACACAACCCAATCCAACCCACCCGATCGCCATAAAGGGTATTTACTGTCGCTGCCGCAGAGCCTCGCTTGACGACTTAACCCAAGCGGTCGTTCCACGTCCATTCTCCGGACGGAGTCAAAGACAAAGGCCGGCGGAGCTGGACAATAGGCAAGGTTGTTGCTTGTGGGTAGAGGGTTTCAATCCCGAAATTGTACCTTTCCCGGGCGAGAAGGGCTTGCATGTGGGCCTTTTCAATTTCGCAGATGAATTTAGATGTAGATGTAGATGCGGATGGGCCGGTCGAGTTAATGCCAATGCAAATTGCGGCGCAATATAACCCAATAATTTGAACCAACTCGCGGAGCAGCGAGGGCATCCTACCCGGTTACCCGGTACTGCATAACAATGAAACGAAACTGGGACAGATCGGTGATGGTTTCTCGCTGTGTGTGCCGTGTTAATCCGTTTGCCATCAGCCAGATTATTAGTCAATTGCAGTCGCAGTCGCAGTTGCAGTTGCAGGGTTTCGCTTTCCTCGTCCTCGTTTCACTTTCGAGTTAGACTTTATTGCAGCATCTTGCAGCAACAATCGGCGCAGTTTGGTAACACGCTGTGCCCTTCCACTTTCCACATTCCACGGCCCAATTCGGCGGATTTAGACGGAATCGAGGGACCCTGACTATGTTCGCATAATGAAAGCGAAACCAAACCGGGTTGCAAAGTCAGGGCATTCCATCGCCGTCGCCATCGCCACCGCCATCTTCTGCGGGCGTTTGTTTGTTTGTTTGCTGGGATTAGCCAAGGGCTTGACTTGGAACCCAATCCCAATCCCAATCCCAATCCCAAAGCCAATCCCAGTGCCCATCCCGATCGCAGTCCCATGCCCTTTTCATTAGAAAGTCATAAAAACACATAATAATGATGTCGAACGGATTAGCTGCGCGCAGTCCAGGCAGCGCAATTAACGGACTAGCAAACTGGGTTATTTCTATTTTTTTATTTTCGCCGACTTAGCCCTGATCCGCGAGCTTAACCCGTTTGAGCCGCAGCAGGTAGCCATCCCCATCCTGGCTTGCGCAAGTGGCAGGTTCGCTGCCCCAGGAGAGCCGTGGAGGCACTCTGGCCACTGGCCTGGTACAGTTGCCGCTGGGCATGATTATATCATCATAATAAATGTTT

>D. erecta eveS2 + 3X Zelda

GAGGCAGGTACTTCCTTTCCAGGTCGCAGATGTAGGTAGATGTCGGCGCAATATAACCCAATAATTTGAACCAACTCGCGGAGCAGCGAGGGCATCCTACCCGGTTACCCGGTACTGCATAACAATGAAACGAAACTGGGACAGATCGGTGATGGTTTCTCGCTGTGTGTGCCGTGTTAATCCGTTTGCCATCAGCCAGATTATTAGTCAATTGCAGTCGCAGTCGCAGTTGCAGTTGCAGGGTTTCGCTTTCCTCGTCCTCGTTTCACTTTCGAGTTAGACTTTATTGCAGCATCTTGCAGCAACAATCGGCGCAGTTTGGTAACACGCTGTGCCCTTCCACTTTCCACATTCCACGGCCCAATTCGGCGGATTTAGACGGAATCGAGGGACCCTGACTATGTTCGCATAATGAAAGCGAAACCAAACCGGGTTGCAAAGTCAGGGCATTCCATCGCCGTCGCCATCGCCACCGCCATCTTCTGCGGGCGTTTGTTTGTTTGTTTGCTGGGATTAGCCAAGGGCTTGACTTGGAACCCAATCCCAATCCCAATCCCAATCCCAAAGCCAATCCCAGTGCCCATCCCGATCGCAGTCCCATGCCCTTTTCATTAGAAAGTCATAAAAACACATAATAATGATGTCGAACGGATTAGCTGCGCGCAGTCCAGGCAGCGCAATTAACGGACTAGCAAACTGGGTTATTTCTATTTTTTTATTTTCGCCGACTTAGCCCTGATCCGCGAGCTTAACCCGTTTGAGCCGCAGCAGGTAGCCATCCCCATCCTGGCTTGCGCAAGTGGCAGGTTCGCTGCCCCAGGAGAGCCGTGGAGGCACTCTGGCCACTGGCCTGGTACAGTTGCCGCTGGGCATGATTATATCATCATAATAAATGTTT

